# Locus coeruleus patterns differentially modulate learning and valence in rat *via* the ventral tegmental area and basolateral amygdala respectively

**DOI:** 10.1101/2020.04.17.047126

**Authors:** Abhinaba Ghosh, Faghihe Massaeli, Kyron D. Power, Tamunotonye Omoluabi, Sarah E. Torraville, Julia B. Pritchett, Tayebeh Sepahvand, Vanessa M. Strong, Camila Reinhardt, Xihua Chen, Gerard M. Martin, Carolyn W. Harley, Qi Yuan

## Abstract

The locus coeruleus (LC), the main source of forebrain norepinephrine, produces phasic and tonic firing patterns that are theorized to have distinct functional consequences. However, how different firing modes affect learning and valence coding of sensory information are unknown. Here bilateral optogenetic activation of rat LC neurons using 10-Hz phasic trains of either 300 msec or 10 sec accelerates acquisition of a food-rewarded similar odor discrimination, but not a dissimilar odor discrimination, consistent with LC-supported enhanced pattern separation and plasticity. Similar odor discrimination learning is impaired by noradrenergic blockade in the piriform cortex (PC). However, here 10-Hz LC phasic light-mediated learning facilitation is prevented by a dopaminergic antagonist in the PC, or by ventral tegmental area (VTA) silencing with lidocaine, suggesting an LC-VTA-PC dopamine circuitry mediates 10-Hz phasic learning facilitation. Tonic stimulation at 10 Hz did not alter odor discrimination acquisition, and was less effective in activating VTA DA neurons. For valence encoding, tonic stimulation at 25 Hz induced freezing, anxiety and conditioned odor aversion, while 10-Hz phasic stimulation produced an odor preference consistent with positive valence. Noradrenergic blockade in the basolateral amygdala (BLA) prevented conditioned odor preference and aversion induced by 10-Hz phasic and 25-Hz tonic light respectively. CTB retro-labeling showed relatively larger engagement of nucleus accumbens projecting neurons over central amygdala projecting neurons in the BLA with 10-Hz LC phasic activation, compared to 25-Hz tonic. These outcomes argue that LC pauses, as well as LC firing frequencies, differentially influence both target networks and behaviour.

## INTRODUCTION

Locus coeruleus (LC) adrenergic neurons fire in both phasic and tonic modes. However, differential behavioural modulation by these contrasting patterns has not been widely studied. Phasic and tonic LC patterns induce waking in mice arguing for similar arousal effects. In sensory coding, tonic patterns enhance thalamic feature selectivity and encoding in rats (1), while phasic LC activation accelerates spatial learning (2) and promotes its consolidation in mice (3). Phasic, not tonic, LC activation enhances salience when phase-locked with sensory stimuli in mice (4).

Here we compare phasic and tonic LC activation on difficult odor discrimination (DOD) learning using similar odor pairs in adult rats. Discrimination of similar odors requires norepinephrine (NE) in the piriform cortex (PC) (5). Blockade of NE input retards DOD learning. We also compare the two patterns on valence. In mice, tonic patterns promote aversions and anxiety (6), but it has not been tested in rats. While phasic LC activation is regarded as learning-promoting, whether it, itself, carries valence is unclear. In neonatal rat odor preference learning, LC-NE activation serves as an unconditioned reward (7). Phasic LC firing to tactile stimuli in neonates mediates preference learning. Tactile stimulation fails to activate LC after the early sensitive period and no longer induces odor preferences (8). Here we test the valence values of LC patterns in adult rats using real-time and conditioned odor preference tests.

## RESULTS

### Validation of light activation of locus coeruleus neurons

Three weeks following LC AAV infusion (for targeting see Supplementary Fig. 1), we observed ChR2 uptake marked by the fluorescence reporter EYFP (Fig. 1A) or mCherry. The transfection rate determined by the overlapping of ChR2 and dopamine beta-hydroxylase (DBH) expressing cells was 73.7 ± 12.8% (n = 8). We conducted *in vivo* optrode LC recordings. Figure 1B shows an LC neuron activated by a 10-sec, 10-Hz light train (30-msec pulses, laser intensity 150 mA). Figure 1C1-C3 shows LC activation by 10-Hz trains with two light intensities and pulse widths. LC firing frequency changes were significantly induced by 10-sec, 10-Hz light at 30-msec duration, 150 mA (F_2,10_ = 8.69, p = 0.006, One-way repeated ANOVA; Fig. 1C1), or 10 Hz at 50-msec duration, 150 mA (F_2,10_ = 25.42, p < 0.001; Fig. 1C2), but not at 30-msec duration, 100 mA (F_2,6_ = 2.48, p = 0.16; Fig. 1C3). The 30-msec, 150-mA light that effectively drove LC firing was used subsequently in recording (Fig. 1D) and behavioural experiments. A 3 (Time) X 5 (Frequency) mixed ANOVA test showed a significant effect of Time (F_2, 48_ = 16.15, p < 0.001). One-way repeated ANOVA revealed that 10-Hz (F_2, 22_ = 30.45, p < 0.001), 15-Hz (F_2, 10_ = 6.16, p = 0.02), and 30-Hz (F_2, 10_ = 8.86, p = 0.006) light significantly increased the firing rate of LC neurons during the light activation. The immediate early gene Npas4 revealed light-induced expression in LC (t = 4.53, p < 0.001; Fig. 1E&F). *In vitro* cell-attached recording shows action currents follow 10 and 20 Hz blue light activation (Supplementary Fig. 2).

**Figure 1.**
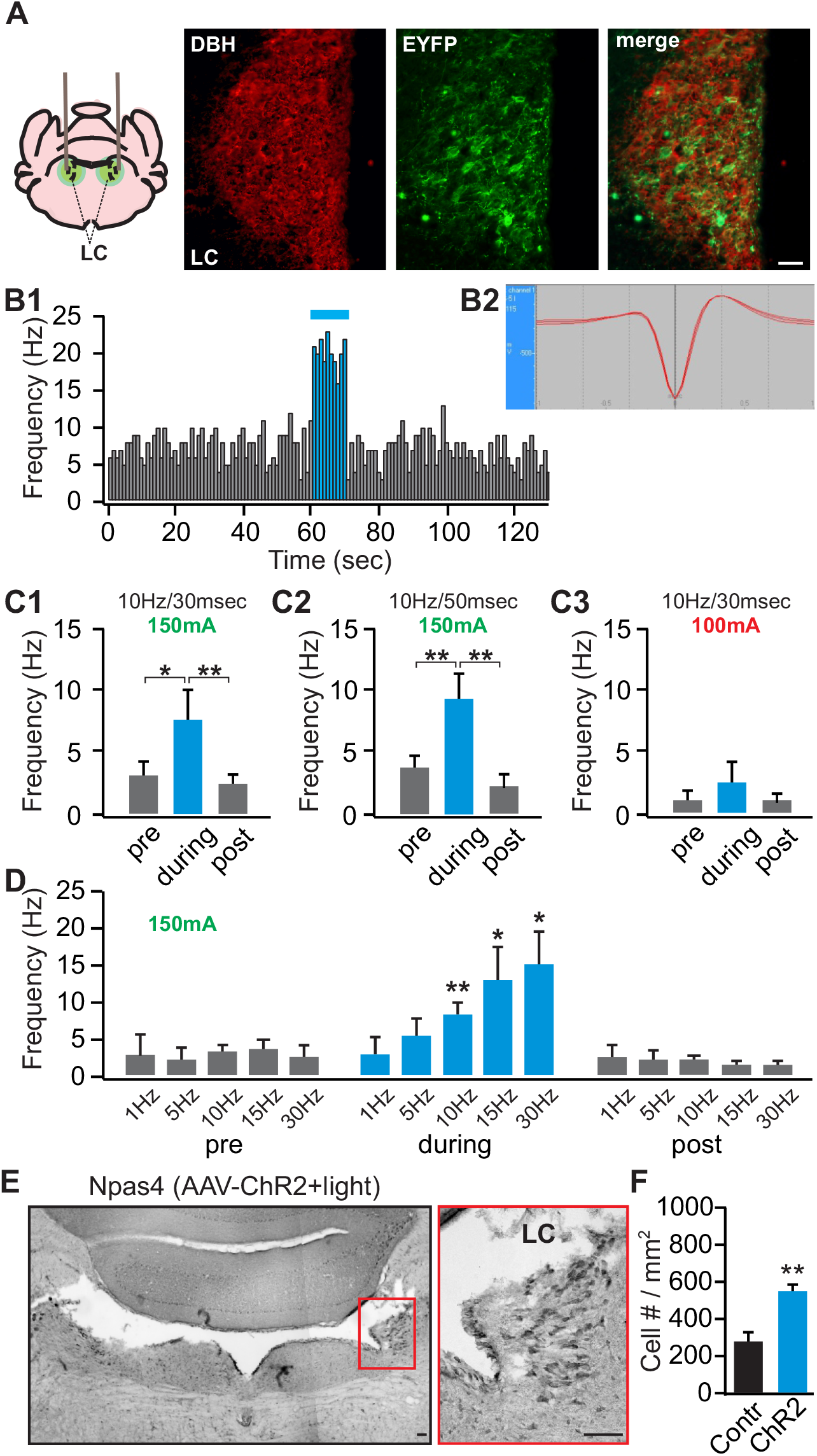
Validation of the light activation of the locus coeruleus (LC) neurons. **A.** Co-localization of DBH (red) and EYFP (green) in the LC of a TH-CRE rat infused with an AAV8-Ef1a-DIO-eChR2(H134R)-EYFP. **B1.** An example of an *in vivo* recording from the LC, showing increased firing of a LC neuron to a 10-Hz, 10-sec light (30-msec duration, 150-mA intensity). **B2.** The waveform of the recorded cell. **C1-C3.** LC firing frequency changes induced by 10-sec, 10-Hz light at 30-msec duration, 150 mA (n = 6, **C1**), at 50-msec duration, 150 mA (n = 6, **C2**), or at 30-msec duration, 100 mA (n = 4, **C3**). **D.** LC responses to light activation with a range of frequencies at 150-mA intensity (n (1/5/10/15/30 Hz) = 2/3/12/6/6). **E.** An example of Npas4 staining of the LC following light stimulation. Right panel is the zoom in of the red square on the left panel. Scale bars, 50 μm. **F.** Npas4^+^ cell counts in the control (n = 6) *vs.* ChR2 rats (n = 7). *p < 0.05; **p < 0.01.

Increasing light frequency elevated LC firing in ~linear fashion in the frequency range <30 Hz (Fig. 1D). While average LC firing rates with 5- and 10-Hz light approximate activation rates, we found LC neurons were not individually driven at those rates *in vivo*. Similar outcomes were observed in mouse *in vivo* where average LC firing with light activation was made up of LC neuron responses of widely varying frequencies (6).

### Different patterns of LC activation show distinct general behavioural effects

We compared 10-Hz phasic (10 sec every 30 sec), 10-Hz tonic and 25-Hz tonic LC activation on open field distance travelled, duration of rearing, and freezing (Fig. 2A-C). For distance traveled in the open field, a 2 (Group) X 3 (Light Pattern) ANOVA showed a significant effect of Light Pattern X Group interaction (F_2, 24_ = 8.831, p = 0.001; Fig. 2A). The ChR2 group showed a reduction in distance traveled with 25-Hz light compared to the control group (t = 4.02, p = 0.002). For duration of rearing with various light patterns, a 2 × 3 ANOVA showed significant effects of Light Pattern (F_2, 24_ = 3.748, p = 0.038), Light Pattern X Group interaction (F_2, 24_ = 13.478, p < 0.001) and Group (F_1, 12_ = 5.356, p = 0.039; Fig. 2B). A one-way repeated ANOVA showed significantly different effects of light patterns in the ChR2 group (F_2, 12_ = 16.783, p < 0.001). Post-hoc Bonferroni tests showed differences between the 10-Hz phasic and 25-Hz tonic (t = 5.745, p < 0.001), and between the 10-Hz tonic and 25-Hz tonic (t = 3.521, p = 0.013). The ChR2 group also showed increased rearing with 10-Hz phasic light compared to the control group (t = 3.910, p = 0.002), and with 10-Hz tonic light (t = 2.181, p = 0.049). For the amount of freezing with various light patterns, a 2 × 3 ANOVA showed significant effects of Light Pattern (F_2, 22_ = 29.050, p < 0.001), Light Pattern X Group interaction (F_2, 22_ = 33.862, p < 0.001), and Group (F_1, 11_ = 31.244, p < 0.001; Fig. 2C). A one-way repeated ANOVA showed significantly different effects of light patterns in the ChR2 group (F_2, 10_ = 36.481, p < 0.001). Post-hoc Bonferroni tests showed differences between the 25-Hz tonic and 10-Hz phasic (t = 7.940, p < 0.001), and between the 25-Hz tonic and 10-Hz tonic (t = 6.697, p < 0.001) activation. The ChR2 group also showed increased freezing with 25-Hz phasic light compared to the control group (t = 6.811, p < 0.001).

**Figure 2.**
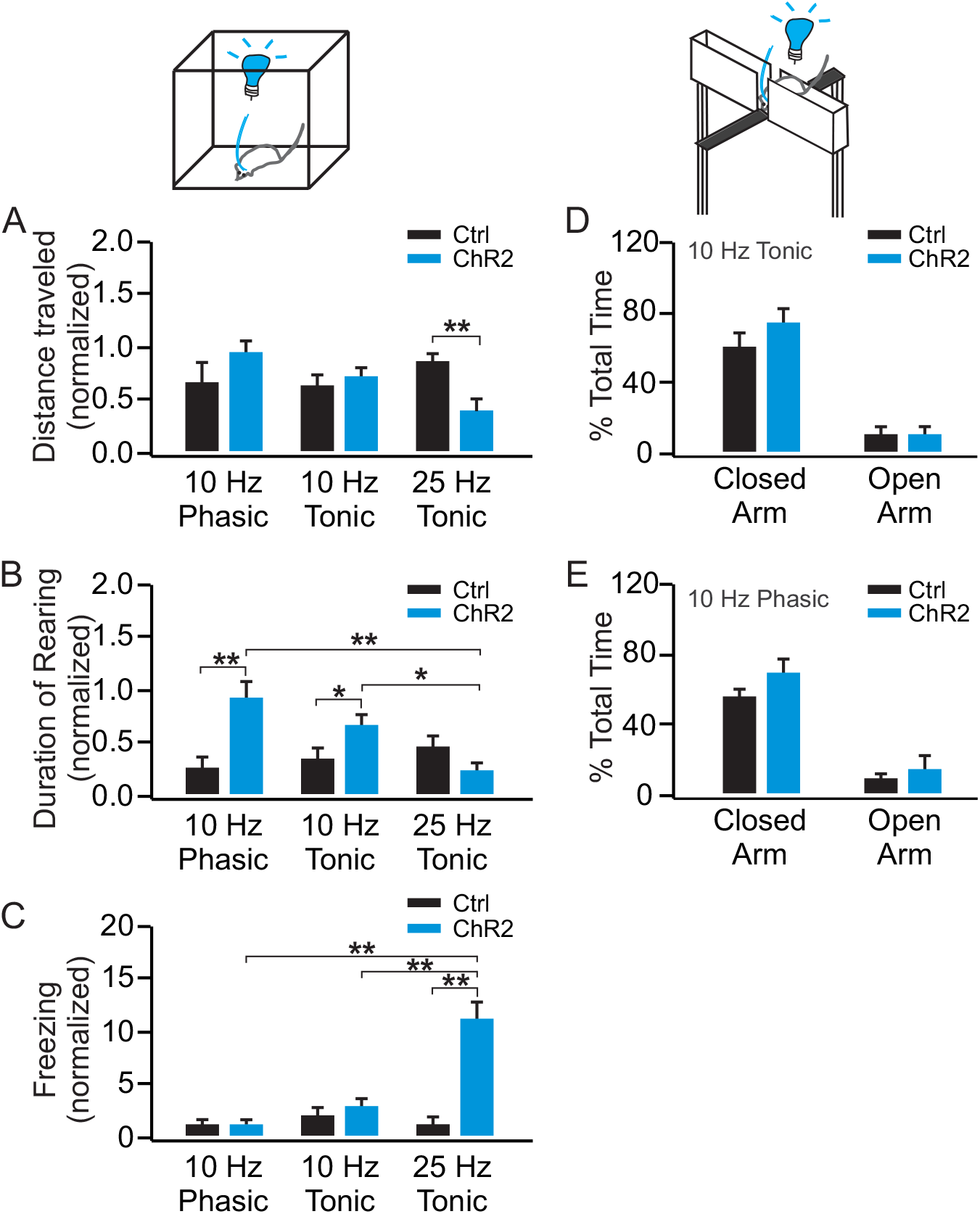
10-Hz phasic LC activation promotes exploration while 25-Hz tonic activation results in increased freezing and decreased mobility. **A.** Distance traveled in the open field with various light patterns in the ChR2 (n = 7) and control (n = 7) rats, normalized to the baseline before the light stimulation. **B.** Duration of rearing with various light patterns in the ChR2 (n = 7) and control (n = 7) rats. **C.** Amount of freezing with various light patterns in the ChR2 (n = 6) and control rats (n = 7). **D.** Percentage time spent in open and close arms of the EPM with 10-Hz tonic light activation in the ChR2 (n = 7) and control (n = 9) rats. **E.** Percentage time spent in open and close arms of the EPM with 10-Hz phasic light activation in the ChR2 (n = 6) and control (n = 6) rats. *p < 0.05; **p < 0.01.

Anxiety during tonic and phasic 10-Hz light patterns was measured in an elevated plus maze (EPM, Fig. 2D & 2E). For percentage time spent in the open and closed arms of the EPM with the 10-Hz tonic light, a 2 (Group) X 2 (Arm) ANOVA showed a significant effect of Arm (F_1, 14_ = 60.312, p < 0.001), however, no effect of Arm X Group interaction (F_1, 14_ = 1.080, p = 0.316), or Group (F_1, 14_ = 2.354, p = 0.147). With 10-Hz phasic light activation, there was a significant Arm effect (F_1, 10_ = 220.25, p < 0.001), but no Arm X Group interaction (F_1, 10_ = 1.135, p = 0.312), or Group effect (F_1, 10_ = 1.657, p = 0.227; Fig 2E).

### LC phasic patterns enhance similar odor discrimination learning

Rats were trained to associate a food pellet with one odor from an odor pair (Fig. 3A). After learning the simple odor discrimination (SOD) with a dissimilar odor pair (Fig. 3B, 3C & 3D), rats learned the DOD with a similar odor pair with LC activation (Fig. 3E, 3F & 3G). Three light patterns were used during DOD training: 10-Hz 10-sec phasic (10 sec every 30 sec; Fig 3B & 3E), 10-Hz brief phasic (300 msec every 2 sec; Fig 3C & 3F) and 10-Hz tonic (Fig 3D & 3G).

**Figure 3.**
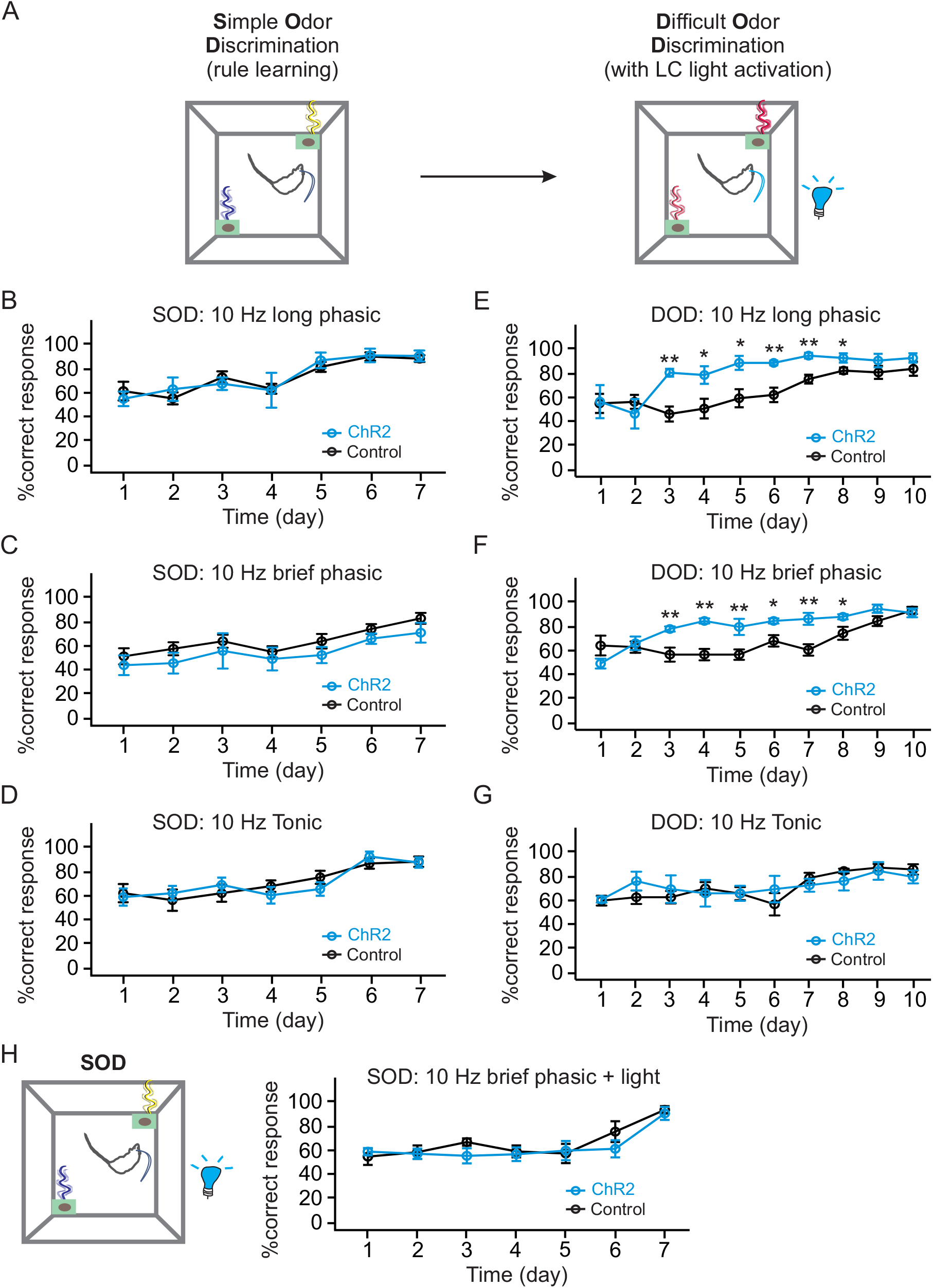
LC phasic patterns, but not tonic pattern, enhance similar odor discrimination learning. **A.** Schematic of odor discrimination learning in rats. Simple odor discrimination (SOD) learning without light is followed by difficult odor discrimination (DOD) learning in the presence of various light patterns. **B.** SOD training in rats with 10-Hz long phasic SOD (n (ChR2/Control) = 5/7). **C.** SOD training in rats with 10-Hz brief phasic SOD (n (ChR2/Control) = 6/8). **D.** SOD training in rats with 10-Hz tonic light (n (ChR2/Control) = 6/7). **E.** DOD training with 10-Hz long phasic. **F.** DOD training with 10-Hz brief phasic. **G.** DOD with 10-Hz tonic light. **H.** SOD training with 10-Hz brief phasic light (n (ChR2/Control) = 6/7). *p < 0.05; **p < 0.01.

ChR2 and control rats showed similar learning in the initial SOD. A two-way mixed ANOVA (Group X Day) showed there was a Day effect (F_6, 60_ = 10.99, p < 0.001 for 10-Hz, 10-sec phasic, Fig. 3B; F_6, 72_ = 9.125, p < 0.001 for 10-Hz brief phasic, Fig. 3C; F_6, 66_ = 12.097, p < 0.001 for 10-Hz tonic light, Fig 3D), but no Group X Day interaction or Group effect in all light patterns. Both brief and longer modes of phasic LC activation accelerated DOD acquisition. For long phasic LC activation (the 10-Hz, 10-sec pattern), there was a significant Day effect (F_9, 90_ = 10.513, p < 0.001), a Day X Group interaction (F_9, 90_ = 2.880, p = 0.005), and a Group effect (F_1, 10_ = 10.069, p = 0.010; Fig 3E). Between group t-tests at each day revealed significantly better correct response rates from day 3-8 with the 10-Hz long phasic pattern (p < 0.05 or p < 0.01). Similarly, for DOD training with 10-Hz brief phasic light (300 msec), there was a significant Day effect (F_9, 108_ = 11.774, p < 0.001), a Day X Group interaction (F_9, 108_ = 4.429, p < 0.001), and a Group effect (F_1, 12_ = 17.47, p = 0.001; Fig 3E). Between group t-tests at each day revealed significantly better correct response rates from day 3-8 10-Hz brief phasic pattern (p < 0.05 or p < 0.01). In contrast, tonic 10-Hz LC activation did not alter acquisition. There was a significant effect of Day (F_9, 99_ = 5.067, p < 0.001), however, no effect of Day X Group interaction (F_9, 99_ = 0.983, p = 0.459), or Group (F_1, 11_ = 0.033, p = 0.860). Linear trend analysis showed improvement with time in both ChR2 (F_1, 5_ = 12.810, p = 0.016) and control groups (F_1, 6_ = 61.491, p < 0.001; Fig 3F).

SOD learning was not affected by phasic 10-Hz LC activation (Fig 3H). A two-way mixed ANOVA revealed a significant effect of Day (F_6, 66_ = 10.713, p < 0.001), but no Group X Day interaction (F_6, 66_ = 0.755, p = 0.608), or Group effect (F_1, 11_ = 0.527, p = 0.483). This underscores NE’s role in discriminations that require pattern separation (5, 9). Enhancement of subtle tactile discriminations by tonic 5-Hz LC activation (1) has also been reported in rats, a frequency not explored here.

Dopamine (DA) co-release from the LC terminals in the hippocampus has been shown to be critical in spatial learning (2) and novelty-mediated memory consolidation (3). We next tested the potential involvement of NE and DA in the PC upon LC phasic light activation in DOD (Fig. 4A). DOD was prevented when the NE antagonists phentolamine and alprenolol were infused in the PC before training, however, D1/5 receptor antagonist infusion selectively abolished LC phasic light induced learning facilitation (Fig. 4B). A mixed ANOVA (Group X Day) showed a significant Day effect (F_9, 207_ = 15.98, p < 0.001), a Group X Day interaction (F_27, 207_ = 5.196, p < 0.001), and a Group effect (F_3, 23_ = 17.73, p < 0.001. *Post-hoc* Bonferroni tests revealed significant differences in learning between the D1/5 antagonist group and ChR2^+^ vehicle group (p = 0.011), and NE antagonist group (p =0.019). There was a significant difference between the NE antagonist group and the ChR2^+^ vehicle group (p < 0.001).

**Figure 4.**
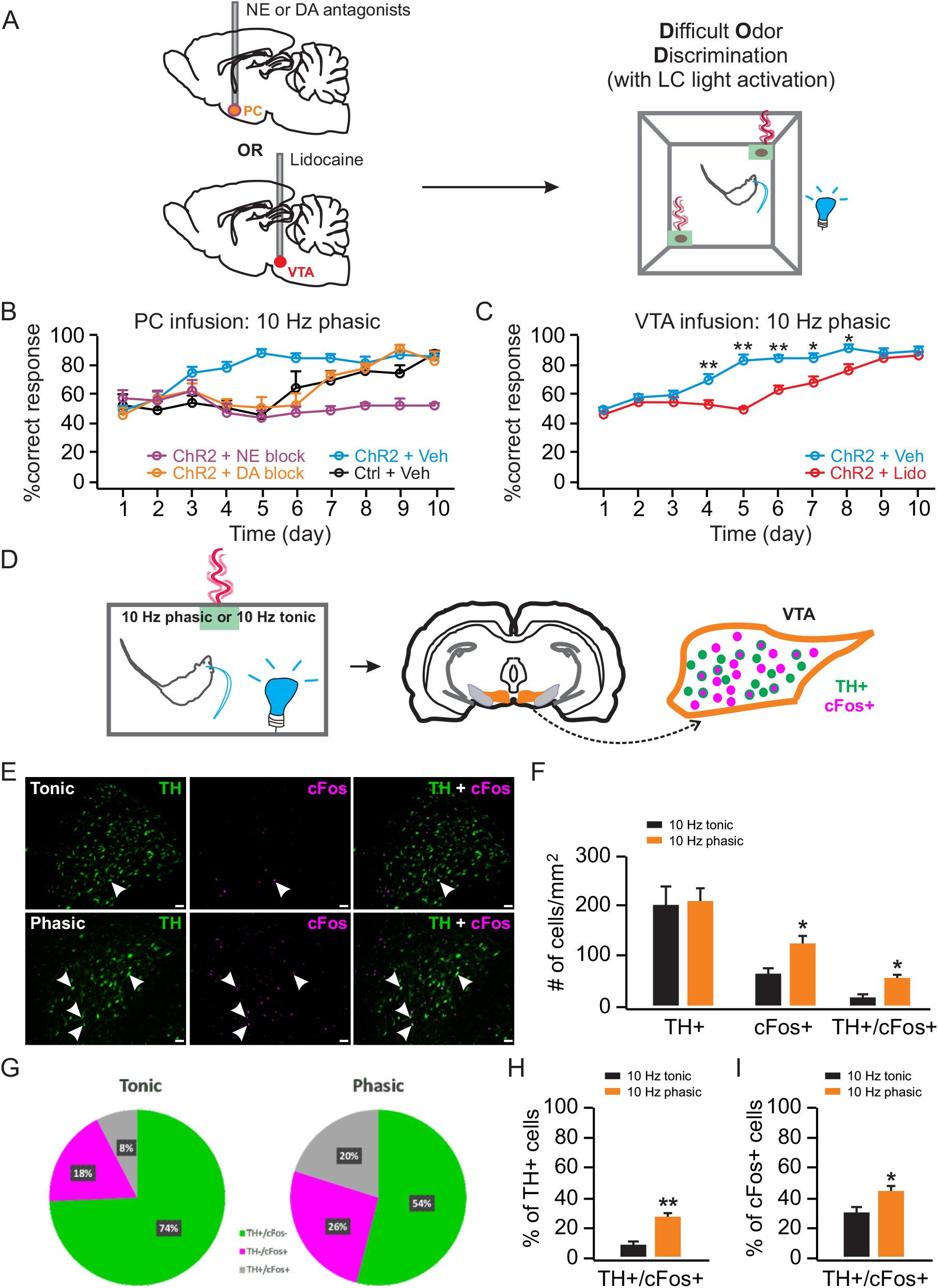
LC phasic activation engages VTA dopamine release to facilitate DOD. **A.** Schematic of DOD training with drug infusion. **B.** DOD training with vehicle or drug infusions in the PC (n (ChR2 vehicle/Control vehicle/ChR2 NE block/ChR2 DA block) = 9/6/6/6). **C.** DOD training with vehicle or lidocaine infusions in the VTA (n (lidocaine/vehicle) = 6/6). **D.** Schematic of measuring cFos expression in the VTA with different LC light patterns. **E.** Examples images of cFos and TH staining in the VTA activated by 10-Hz tonic (upper panels) and 10-Hz phasic light (lower panels). Arrows indicated example TH^+^/cFos^+^ cells. Scale bars, 50 μm. **F.** Total cFos^+^, TH^+^ and TH^+^/cFos^+^ cells activated by 10-Hz tonic and phasic lights (n (tonic/phasic) = 3/4). **G.** Percentage activation of TH^+^ and cFos^+^ cells. **H.** Percentage TH^+^/cFos^+^ cells over total TH^+^ population. **I.** Percentage TH^+^/cFos^+^ cells over total cFos^+^ population.*p < 0.05; **p < 0.01.

These results argue that while NE is essential for pattern separation-dependent odor discrimination to occur, DA release in the PC during LC phasic light mediates the learning facilitating effect. There are two possible scenarios: either LC axon terminals co-release DA upon LC phasic activation (2, 3), or VTA releases DA into the PC (10, 11) upon LC activation. To determine the source of DA released during odor discrimination learning, we infused lidocaine into the VTA to silence the VTA during DOD. Lidocaine infusion prevented the learning facilitation effects of the LC phasic light (Fig. 4C), but not the acquisition of DOD learning. A mixed ANOVA (Group X Day) showed a significant Day effect (F_9, 90_ = 41.46, p < 0.001), a Group X Day interaction (F_9, 90_ = 5.60, p < 0.001), and a Group effect (F_1, 10_ = 90.94, p < 0.001). T-tests showed significantly higher correct response rates of the ChR2^+^ vehicle group relative to the ChR2^+^ lidocaine group on days 4-8 (P < 0.05 or P < 0.01).

Why does phasic LC activation promote DOD while tonic activation does not? In other words, does LC phasic activation engage DA neurons in the VTA more effectively than tonic activation? We next studied neuronal activation patterns in the VTA by phasic *vs* tonic LC activations. We measured cFos expression in the VTA following either 10-Hz phasic or tonic LC activation paired with an odor (Fig. 4D). The 10-Hz phasic pattern activated significantly more cFos^+^ cells in the VTA (t = 3.778, p = 0.012) and had a larger number of TH^+^/cFos^+^ cells compared (t = 3.781, p = 0.013) than that of 10-Hz tonic activation (Fig. 4E and 4F). Twenty percent of the cells measured were both TH^+^ and cFos^+^ with the LC phasic activation, in contrast to 8% in the tonic activation condition (Fig. 4G). The percentage of cFos^+^ cells in the total TH^+^ population is also significantly higher with the 10-Hz phasic pattern than with the 10-Hz tonic pattern (t = 6.199, p = 0.002; Fig. 4H). The percentage of TH^+^/cFos^+^ cells over total cFos^+^ cells is also higher in the 10-Hz phasic group (t = 2.912, p = 0.033; Fig. 4I). These results argue that the 10-Hz LC phasic pattern engages VTA DA neurons more effectively than 10-Hz tonic. This may explain the learning facilitation effect of PC DA with phasic, but not tonic, LC activation.

### LC phasic and tonic patterns promote differential odor valence learning

We tested phasic and tonic LC light patterns in a real time odor preference test (ROPT) and a conditioned odor preference test (COPT; Fig 5A). In the ROPT, rats were tested for the time spent in the two odor zones, first in the absence, then in the presence, of light activation associated with one odor (O1). Ten-Hz phasic activation increased the time rats spent in the light-paired odor zone (Fig. 5B). A 2 × 2 × 2 (Group X Odor X Time) ANOVA showed a significant effect on the Group X Odor X Time interaction (F_1, 15_ = 5.529, p = 0.033). Follow up 2 × 2 ANOVAs showed a significant Odor X Time interaction in the ChR2 group (F_1, 8_ = 8.266, p = 0.021), but not in the control group (F_1, 7_ = 0.80, p = 0.786). ChR2 rats spent significantly more time in O1 during the 10-Hz phasic light, compared to the baseline no light condition (t = 3.64, p = 0.007). On the contrary, 10-Hz tonic light had no significant effect in the Group X Odor X Time interaction in ROPT (F_1, 15_ = 4.221, p = 0.06), and there was no group difference (F_1, 15_ = 0.906, p = 0.356; Fig. 5C).

**Figure 5.**
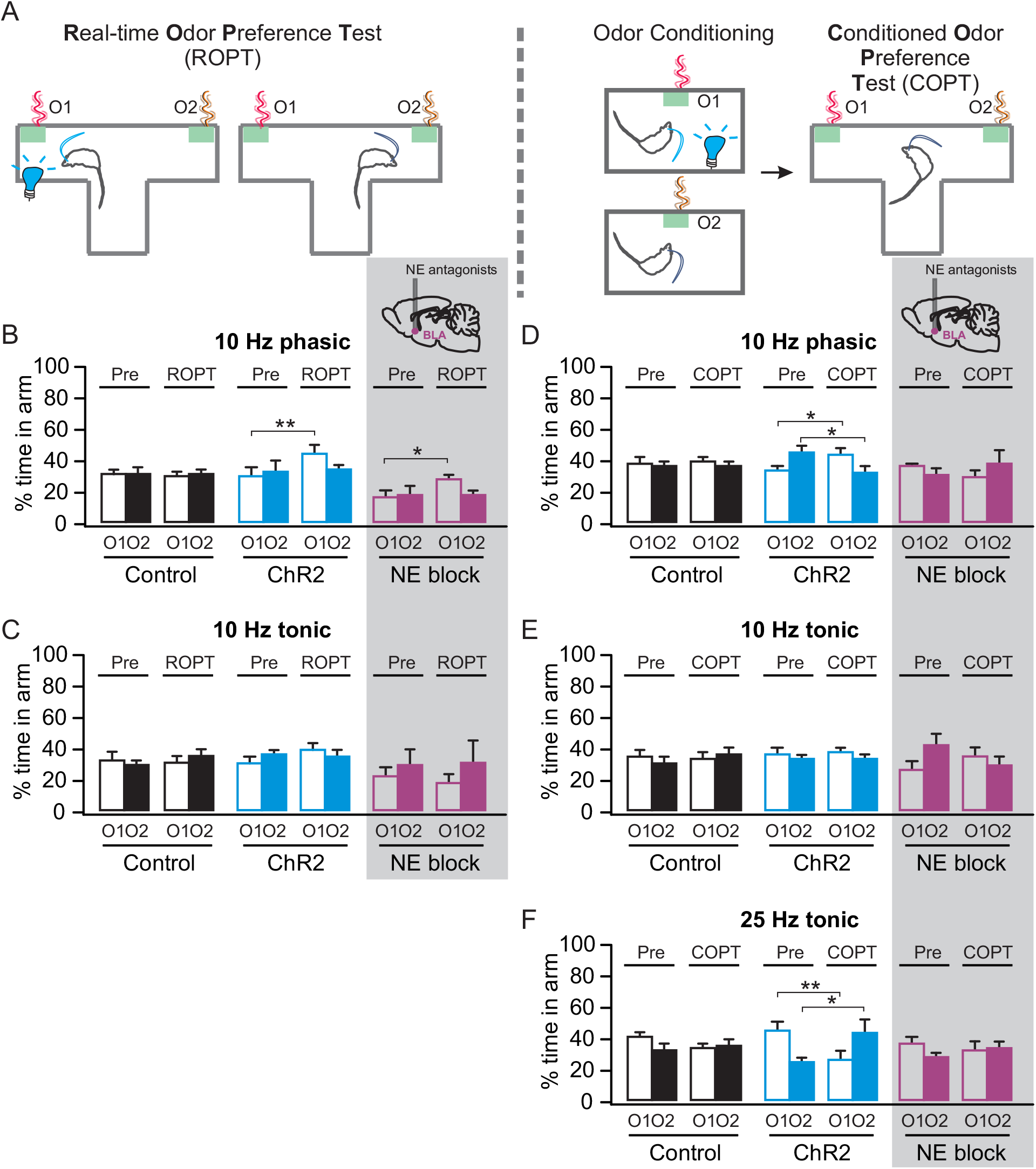
Twenty five-Hz tonic LC activation leads to conditioned odor aversion while 10-Hz phasic pattern results in odor preference. **A.** Schematic of real-time odor preference test (ROPT) and conditioned odor preference test (COPT). **B.** Percentage time spent in each odors in ROPT, with 10-Hz phasic light paired with O1 (n (ChR2/Control/NE block) = 9/8/7). **C.** Percentage time spent in each odor in ROPT, with 10-Hz tonic light paired with O1 (n (ChR2/Control/NE block) = 10/7/5). **D.** Percentage time spent in each odor in COPT, with 10-Hz phasic light conditioned with O1 (n (ChR2/Control/NE block) = 11/12/6). **E.** Percentage time spent in each odor in COPT, with 10-Hz tonic light conditioned with O1 (n (ChR2/Control/NE block) = 11/10/5). **F**. COPT with 25-Hz tonic light (n (ChR2/Control/NE block) = 11/8/5). *p < 0.05; **p < 0.01.

In the COPT, rats were light-stimulated in the presence of one odor (O1). Time spent with O1 *vs*. a control odor O2, before and after conditioning was compared. There was a conditioned preference with 10-Hz phasic, but not 10-Hz tonic pairing. With 10-Hz phasic light conditioned with O1, there was a significant Group X Odor X Time effect (F_1, 21_ = 4.562, p = 0.045; Fig. 5D). Follow up 2 × 2 ANOVAs showed a significant Odor X Time interaction in the ChR2 group (F_1, 10_ = 8.212, p = 0.017), but not in the control group (F_1, 11_ = 0.034, p = 0.857). ChR2 rats spent significantly more time in O1 (t = 2.25, p = 0.048) and less time in O2 (t = 2.41, p = 0.037) after conditioning with the 10-Hz phasic light. However, with 10-Hz tonic light paired with O1, there was no significant Group X Odor X Time conditioning effect (F_1, 19_ = 0.904, p = 0.354; Fig. 5E).

The immobility associated with the tonic 25-Hz light (Fig. 2C) excluded ROPT testing. In COPT, we found conditioned avoidance of the 25-Hz light-paired odor (Fig. 5F). A 2 × 2 × 2 (Group X Odor X Time) ANOVA showed significant effects of the Group X Odor X Time interaction (F_1, 14_ = 5.797, p = 0.030), and Groups (F_1, 14_ = 7.923, p = 0.014). Follow up 2 × 2 ANOVAs showed a significant Odor X Time interaction in the ChR2 group (F_1, 14_ = 14.971, p = 0.002). ChR2 rats spent significantly less time in the O1 arm after it was associated with the 25-Hz tonic light, compared to the baseline before odor-light conditioning (t = 3.86, p = 0.006), while they spent more time in the control odor O2 (t = 3.77, p = 0.007). This replicates the conditioned place aversion seen with tonic 5-Hz activation in mice (6).

In mice, it has been shown that LC tonic activity promotes aversive behavior through basolateral amygdala (BLA) β-adrenoceptors (12). The BLA is a critical site for valence associative learning (13). BLA has been found to contain functionally distinct neuronal populations projecting to either negative-(central amygdala, CeA) or positive-(nucleus accumbens, NAc) valence encoding circuitry (14, 15). It is plausible that the LC-BLA projection is involved in the odor valence encoding observed here with differential LC light activation patterns. Here we first tested the involvement of BLA adrenoceptors in COPT with either phasic or tonic LC activation. BLA α- and β-noradrenergic blockade with phentolamine and alprenolol prevented both LC 10-Hz phasic light induced odor preference and LC 25-Hz tonic induced odor aversion. When comparing the effect of 10-Hz phasic light on the NE blockade group with the control and ChR2 non-infusion groups, a 2 × 2 × 3 (Time X Odor X Group) ANOVA showed a significant Time X Odor X Group effect (F_2,26_ = 5.257, p = 0.012; Fig. 5D). A follow up ANOVA with the NE blockade group showed no significant effects of Time (F_1,5_ = 0.02, p = 0.967), Odor (F_1,5_ = 0.167, p = 0.700), or Time X Odor interactions (F_1,5_ = 2.778, p = 0.156). Similarly, the 10-Hz tonic light did not produce any odor valence effects in COPT in the NE blocker group (Time X Odor interactions: F_1,4_ = 2.775, p = 1.717; Fig. 5E). With 25-Hz tonic light, a 2 × 2 × 3 (Time X Odor X Group) ANOVA showed a significant Time X Odor X Group effect (F_2,23_ = 3.873, p = 0.036; Fig. 5F). A follow up ANOVA with the NE blockade group showed no significant effects of Time (F_1,4_ = 0.090, p = 0.779), Odor (F_1,4_ = 1.407, p = 0.301), or Time X Odor interactions (F_1,4_ = 2.775, p = 0.171), arguing that the aversive odor conditioning by 25-Hz light was mediated through BLA adrenoceptors. The observed effect is not due to cannular infusion, as vehicle infusion with 25-Hz light activation produced a similar odor aversion as in the ChR2, non-infused group (Supplementary Fig. 3). Additionally, NE blockade did not prevent the ROPT odor preference observed with 10-Hz phasic light (F_1,6_ = 9.688, p = 0.021). The NE blockade group still spent significantly more time in O1 in the presence of the 10 Hz-light in ROPT (t = 2.581, p = 0.041; Fig. 5B). Therefore, while BLA NE mediates conditioned valence learning dependent on differential patterns of LC activation, this circuitry is not involved in real-time preference in ROPT. Phasic light mediated enhanced exploration (Fig. 2B) may explain the acute effect of 10-Hz phasic light.

After establishing the requirement of BLA NE in LC light mediated valence learning in COPT, we next tested whether tonic and phasic activation of the LC biases activation of the BLA ensembles projecting to aversive and reward circuitry respectively. We infused retro-tracing dyes linked to Cholera Toxin B (CTB) in the central amygdala (CeA) and nucleus accumbens (NAc), and examined the overlap of CeA or NAc projecting neurons with cFos^+^ cells in the BLA activated by either 10-Hz phasic or 25-Hz tonic LC lights (Fig. 6A and 6B). The CTB labeled CeA (t = 1.695, p = 0.165) and NAc (t = 0.658, p = 0.546) projecting cell numbers were comparable in the two groups (Fig. 6C). Although the two LC light patterns activated similar numbers of cFos^+^ cells in the BLA (t = 0.658, p = 0.546; Fig. 6C), the distribution patterns of cFos^+^ cells were dramatically different (see example images in Fig 6B). With 25-Hz tonic light, 37% of the cFos^+^ cells were CeA projecting, while only 9% were NAc projecting (Fig. 6D). In contrast, 31% cFos^+^ cells were NAc projecting and only 5% were CeA projecting with the 10-Hz phasic pattern (Fig. 6D). The percentage of cFos^+^ cells among CeA projecting cells is significantly higher in the 25-Hz tonic group (t = 4.310, p = 0.013; Fig. 6E), whereas the percentage of cFos^+^ cells among NAc projecting cells is greater with the 10-Hz phasic pattern (t = 3.765, p = 0.020; Fig. 6F). Taken together, the 10-Hz LC phasic pattern preferentially activates NAc projecting neurons in the BLA, whereas the 25-Hz tonic LC activation preferentially engages CeA projecting neurons.

**Figure 6.**
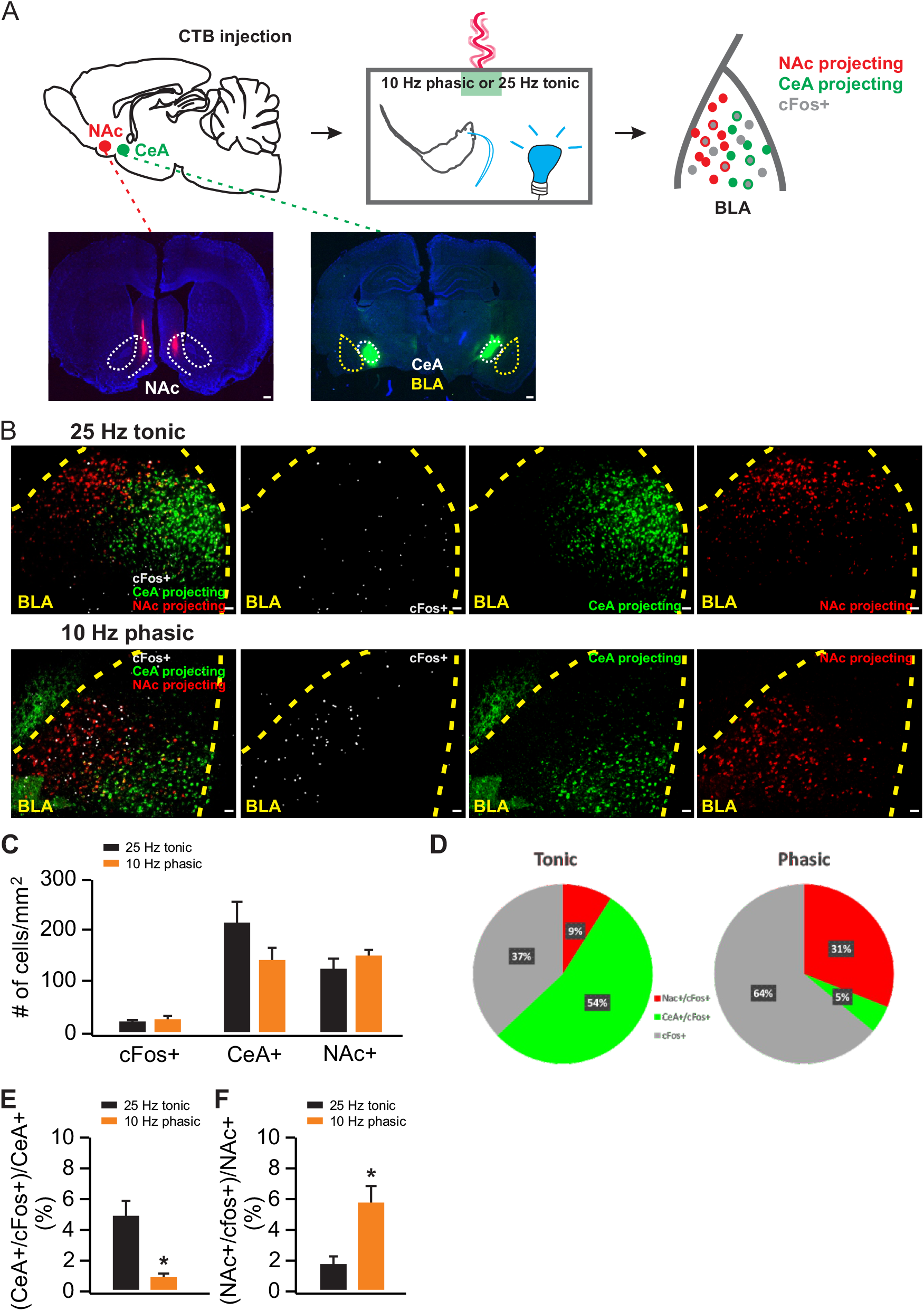
Ten-Hz phasic and 25-Hz tonic LC activation engage positive and negative projecting circuitry respectively in the BLA. **A.** Schematic of measuring cFos activation in the BLA with CTB labeling NAc and CeA projecting neurons. **B.** Examples images of cFos, CTB-488 (labeling CeA projecting neurons) and CTB-594 (labeling NAc projecting neurons) in the BLA activated by 25-Hz tonic (upper panels) and 10-Hz phasic light (lower panels). Scale bars, 50 μm. **C.** Total cFos^+^, CeA^+^ and NAc^+^ cells activated by tonic and phasic lights (n (tonic/phasic) = 3/3). **D.** Percentage activation of cFos^+^, CeA^+^ and NAc^+^ cells. **H.** Percentage CeA^+^/cFos^+^ cells over total CeA^+^ population. **I.** Percentage NAc^+^/cFos^+^ cells over total NAc^+^ population. *p < 0.05.

## DISCUSSION

This work aimed to further our understanding of whether and how the LC produces firing mode-specific behavioural responses. We assessed how LC activation patterns differentially modulate general behaviour, odor discrimination learning and valence encoding. In general behaviour, our assessments revealed increased exploration with both 10-Hz phasic and tonic activation expressed as increased rearing. With 25-Hz tonic activation there was reduced distance traveled and increased freezing. An increase in rearing has not been previously reported with LC activation, but hippocampal NE infusions increase open field rearing (16, 17) as seen here. Freezing with 25-Hz tonic LC activation is similar to the report of behavioural arrest with tonic LC activation (10-25 Hz) in mice (18). However, in the mice, 3-Hz tonic activation also increased distance traveled, while 10-Hz phasic activation decreased it. In rats, neither distance traveled in the open field nor anxiety in the EPM, was altered by phasic or tonic 10-Hz LC activation. The rat outcome also contrasts with mice that show increased anxiety to 5-10 Hz LC tonic activation as indexed by decreased center open field entries and increased closed arm time in an EPM (6).

Anxiety differences between mice and rats with 10-Hz LC activation may relate to species differences or to total LC-NE release, which could be less in rat as fewer cells are likely to be recruited given LC size relative to the optic fiber. Behavioural inhibition in mice was linked to a decrease in NE release as indexed by micro-dialysis during 10 min of a 10-Hz LC light (18). Presynaptic NE exhaustion was proposed to explain the NE reduction, however, we have also documented inhibition of LC after 2 min of 10-Hz light activation (19).

These LC effects argue for enhanced attention or plasticity with phasic LC activation accelerating pattern separation learning here, consistent with previous reports of accelerated spatial learning (2) and perceptual learning (20) with optogenetic phasic LC activation. Since these tasks were multi-trial, enhancement of plasticity-related proteins by phasic activation, previously shown to promote consolidation (20), could also contribute to faster acquisition.

A theory of LC-NE function based on monkey data (21) proposes that phasic firing facilitates performance and optimizes behavior during focused task performance (exploitation), while tonic firing interferes with phasic coding and facilitates exploration (21). Optogenetic studies mimicking these conditions, including the present study, are consistent with a role for phasic LC activation in enhancing task performance. We show phasic LC activation accelerates learning even when not time-locked to stimuli or to decisions. In contrast to the exploit/explore hypothesis, however, phasic LC activation also promoted exploration. Only 25-Hz tonic activation had negative valence associations and inhibited exploration. This high tonic aversive result is congruent with reports of freezing, anxiety and conditioned avoidance with lower tonic frequencies in mice (12). Here phasic and tonic stimuli were both 10 Hz. While higher NE release with phasic activation is explained as due to a higher phasic frequency (22), here such an explanation is unlikely. In the present study, cumulative release should be higher with 10-Hz tonic than 10-Hz phasic patterns since frequency is constant. Single SOD trials are completed rapidly making it unlikely that transmitter exhaustion occurred.

Hippocampal-dependent learning has been shown to involve LC co-released DA (2, 3). DA release from the LC axonal terminals in the dorsal hippocampus, but not NE, mediates spatial learning (2) and novelty mediated learning enhancement (3). Whether co-release of DA from LC axons exists in other structures is not known. Here we show that while DA mediates the facilitating effect of odor discrimination with the 10-Hz phasic LC activation, the effect is mediated by an indirect circuitry *via* VTA activation by the LC phasic light and release of DA in the PC. This highlights structure-dependent mechanisms of LC release and modes of actions. In odor valence conditioning, however, the valence encoding appears to be directly mediated by NE release to the BLA. Noradrenergic antagonist infusion in the BLA prevented both LC 10-Hz phasic induced preference and 25-Hz tonic light induced avoidance.

Recent evidence suggests that functional heterogeneity within the LC itself is responsible for the distinct roles of the LC (23–26). However, it is also possible that heterogeneity of release effects within LC projection sites also plays a key role in the functional diversity of the LC. Our results demonstrate that indeed this is the case. We have shown that 10-Hz phasic LC activation preferentially engages VTA DA neurons compared to a 10-Hz tonic light. This appears to explain why 10-Hz phasic LC activation is more effective in promoting accelerated odor discrimination learning. We have also demonstrated that phasic and tonic patterns, which lead to differential valence outcomes in our study, engage different neuronal ensembles in the BLA. The BLA, a projection site of the LC, has been found to contain functionally distinct neuronal populations projecting to either negative-(CeA) or positive-(NAc) valence encoding circuitry (14, 15). We show that tonic and phasic activation of the LC biases activation towards neuronal ensembles in the BLA projecting to aversive and reward circuitry respectively, thereby highlighting the role of downstream projection site heterogeneity in differential LC functions.

We, and others, have suggested higher NE concentrations promote plasticity because they engage low affinity β-adrenoceptors linked to the cAMP/PKA/CREB cascade (27), but simple concentration effects may not differentiate tonic and phasic effects here. It appears that the pause associated with the phasic pattern is, in some way, essential for learning acceleration. A similar conclusion is reached when examining DA’s role in providing a teaching signal (28). Differential engagement of neuronal ensembles with noradrenergic receptor heterogeneity likely plays a critical role in LC firing-mode dependent behavioural effects.

In summary, the present outcomes support Aston-Jones and Cohen’s hypothesis (21) that LC phasic bursts promote optimized adaptation as proposed, but phasic patterns can also generate exploration. Frequency is critical for tonic effects with a report of discrimination benefits at 5 Hz in rats (1) and aversive valence at 5 Hz in mice (6). The present experiments reveal aversive valence at 25-Hz, but not 10-Hz tonic activation, in rats, and, for the first time, demonstrate positive valence with phasic 10 Hz optogenetic LC activation in adult rat.

The several network mechanisms by which differing patterns and frequencies of LC activation achieve specific behavioural outcomes revealed in these experiments means that there is still a great deal of work remaining to understand LC’s functional contributions. Differing receptor populations in target structures, differing concentrations of NE release, effects of co-release of peptides and amino acids, and the conditions under which they occur are all open questions for future investigations. The physiological mechanisms illuminated in these experiments will need to be integrated with the recent structural evidence of greater specificity of LC subpopulation targeting than previously suspected (23, 25). The interaction of these more selective physiological and structural features of LC modulation of brain networks promises a broad array of insights to come with novel functional implications for both basic and clinical brain science.

## MATERIAL AND METHODOLOGY

### Animals and Ethics Statement

TH-CRE homozygous male breeders (Sage laboratories, Boyertown, PA) were bred with Sprague-Dawley (SD) female breeder rats (Charles River, Saint-Constant, Canada) for TH-CRE heterozygous offspring that were used in this study. Rats of both sexes were housed in 12 h light/dark cycle and had *ad libitum* access to food and water unless during food deprivation for experiments. During food deprivation, each rat was given 20 g of regular rat chow/day and was monitored for body-weight and health status on weekly basis. All experimental protocols followed the guidelines of Canadian Council of Animal Care and were approved by Memorial University animal care committee.

### Viral Transduction

Adeno-associated virus (AAVdj or AAV8) served as a vector to carry the key construct of channelrhodopsin 2 (ChR2) with reporter gene for fluorescent proteins (EYFP or mCherry) under a double-floxed inverted open reading frame (DIO). Experimental constructs were AAVdj-EF1a-DIO-hChR2(H134R)-mCherry or AAV8-Ef1a-DIO-eChR2 (H134R)-EYFP. Control construct was AAVdj-EF1a-DIO-mCherry. All AAVs were provided by Deisseroth Laboratory at Stanford University.

### Stereotaxic Surgery

Three to five months old adult TH-Cre rats received bilateral virus infusion (5e12/ml) in LC under isofluorane anesthesia in a stereotaxic frame. Each hemisphere received two infusions, each of 0.7 μl (fluorescent beads: virus = 2:5) at the rate of 0.5 μl/min. The cannula was lowered at a 20° angle to avoid transverse sinus. Infusion coordinates were 11.8-12.2 mm posterior, 1.2 and 1.4 mm bilateral, and 6.3 mm ventral with respect to bregma. Minimum one month after infusion surgery, rats underwent cannula (ferrule attached, containing optical fiber; Doric Lenses, Quebec, Canada) implantation surgery (11.8-12.2 mm posterior, 1.3 mm bilateral, and 6.3 mm ventral with respect to bregma), followed by minimum two weeks of recovery period before commencing behavioural tests.

For experiments requiring drug infusion, metal infusion cannulas were implanted in BLA (AP: 2.5 mm posterior, ML: 4.9mm bilateral, DV: 7.8 mm) (29), or PC (AP: 1.8-2.0 mm anterior, ML: 4 mm bilateral, DV: 7.3-7.4 mm) (5), or VTA (AP: 5.3 mm posterior, ML: 1mm bilateral, DV: 8.1 mm) (30, 31) combined with LC optical fiber cannula implantation.

For experiments requiring CTB infusions, surgeries were combined with LC optical fiber cannula implantation and were allowed 10 days recovery before carrying out experiments. CTB-594 and CTB-488 (1% w/v in phosphate buffer; Invitrogen, Waltham, MA) were infused by separate 32g bevelled 1μl Hamilton syringes (Neuros 7001 KH) attached to an infusion pump (Pump 11 Elite; Harvard Apparatus, Saint-Laurent, Canada) (12, 32) in NAc (200 nl; AP: 1 mm anterior, ML: 1 mm bilateral, DV: 6.5 mm) and CeA (150 nl; AP: 2.1 mm posterior, ML: 4.2 mm bilateral, DV: 7.5 mm) respectively.

### *In vivo* electrophysiology

Rats underwent *in vivo* electrophysiology experiment one month post-infusion. Rats were anaesthetized with 15% urethane at 10 ml/kg of body weight and placed in a stereotaxic frame. Surgical procedure was carried out following appropriate sterile methods. A hole was drilled in the skull (12.3 mm posterior, 1.3 mm left lateral to bregma) and an optrode, assembled just before the experiment (400 μm glass optical fiber; Thorlabs Inc, Newton, NJ), bundled with 200/280 μm tungsten electrode; FHC, Bowdoin, ME), was lowered down at 20° angle to 6.1-6.9 mm ventral to brain surface until LC neurons were identified by slow spontaneous spiking and burst response to toe pinch (audio-monitor and oscilloscope response) (19). Glass optical fiber was connected to a Laser diode fiber light source (Doric Lenses, Quebec, Canada) by mono-fiberoptic patch cord (0.48 NA, 400/430 μm). Blue light of 450 nm at a power of 90 mW for 400 μm core and 0.48 nA optical fiber was applied. Light pattern was controlled from a Doric software.

Data were acquired and analyzed by SciWorks (DataWave/A-M Systems, Sequim, WA). Signal was detected as a lowest threshold of 1.5X amplitude of the background. Autosort protocol was used to isolate similar cellular waveforms and cluster them in a cell-specific manner. Clusters only with LC-spike characteristics were further analyzed.

### Behavioural tests

#### Light stimulation for behavioural experiments

Two to four weeks after cannula implantation surgery, rats underwent behavioural tests. Bilateral photostimulation at 450 nm – 90 mW was carried out by two Laser light sources (LDFLS_450; Doric Lenses, Quebec, Canada) through mono fiberoptic patch cords. Current equivalence of the power was 150 mA. A total of 4 different photo stimulation patterns were used in the behavioural tests, namely: 10-Hz long phasic (10 Hz, 10 sec every 30 sec); 10-Hz brief phasic (3 pulses every 2 sec); 10-Hz tonic; and 25-Hz tonic. Different patterns of stimulation were controlled from Doric software. Behavioural sessions were video-recorded by ANY-Maze software (Stoelting, Wood Dale, IL) and analyzed offline. Subsets of experiments were analyzed by persons who were blind to the experimental conditions.

### Drug infusion

Drugs or vehicle were infused 30 min before behavioral testing through a 10 μl Hamilton syringe and infusion pump. Lidocaine (4%; Sigma, Oakville, Canada) was infused in VTA (0.3 μl/ hemisphere over 3 minutes with an additional minute before withdrawing the syringe) (31). For PC infusions, 1 μl drug or vehicle was infused per hemisphere over 2 min followed by 1 min before withdrawing the syringe. D1/D5 receptor antagonist SCH 23390 (3.47 mM, Sigma, Oakville, Canada), α-adrenoceptor antagonist phentolamine hydrochloride (10 mM; Sigma, Oakville, Canada) and β-adrenoceptor antagonist alprenolol hydrochloride (120 mM; Tocris Bioscience, Bristol, UK) were used for PC infusions (2, 5).

#### General Behavioural Experiments

##### Open Field Maze

Rats explored an opaque plexiglass box (60 cm × 60 cm × 40.5 cm) with black bottom for a 10 min session daily for four days consecutively while receiving no stimulation on day 1, 10-Hz long phasic stimulation on day 2, 10-Hz tonic stimulation on day 3 and 25-Hz tonic stimulation on day 4. During each session, distance travelled, time spent rearing (both free and supported) and time spent freezing were recorded and analyzed.

##### Elevated Plus Maze

Following a 15-min photo-stimulation in the homecage, rats were placed at the central area of the elevated plus maze (50 cm × 10 cm each arm, 38 cm wall on the closed arms, 11 cm × 11 cm central platform, 52 cm high from the ground, inside painted black) facing the open arm opposite to the experimenter. Then rats spent 5 min in the EPM while the stimulation continued. Time spent in closed and open arms was counted and recorded.

#### Olfactory Behavioural Experiments

##### Odorants

Odors used in the experiments were listed in Table 1.

| Name of the test | Odor 1 (O1; *\*associated with photostimulation*) | Odor 2 (O2) |
| --- | --- | --- |
| Dissimilar odor discrimination | Almond extract | Coconut extract |
| Similar odor discrimination | Heptanol and Octanol (40:60 mixture; 0.001%) | Heptanol and octanol (50:50 mixture; 0.001%) |
| Valence test (10 Hz long phasic) | Vanilla (2%) | Peppermint (2%) |
| Valence test (10 Hz tonic) | Orange (2%) | Propanoic acid (0.033%) |
| Valence test (25 Hz tonic) | Benzaldehyde (0.05%) | Isoamyl acetate (0.05%) |

The odor concentrations for similar odor discrimination and valence tests were chosen either based on previous publication (5) or estimated vapor pressure of 1 Pascal (33).

##### Odor Discrimination Learning

Rats were food deprived for 4-7 days before the onset of the experiments and food deprivation continued during the course of the experiment. Five days of habituation for context (60 cm × 60 cm × 40.5 cm box), sponge and positive reinforcement (chocolate cereal) were conducted first. Afterwards rats performed odor discrimination task 10 trails/day, each trial being maximum 3-min long. Two sponges containing odor 1 (O1) and odor 2 (O2) were presented randomly in two corners of the box. The O1 sponge had a 2 cm^2^ hole at the top center, containing a retrievable chocolate cereal. To balance the smell of the chocolate cereal, a non-retrievable chocolate cereal was placed in a hidden hole in the O2 sponge. Between trials rats were confined to a “home corner” in the box with an L-shaped plexiglass barrier for 20 sec while sponge positions were changed. During trials rats were allowed to explore the box and sniff the sponges. Trials ended as soon as it made a nose-poke inside the hole, irrespective of the odor identity. Response was considered correct if nose-poke was in O1 sponge. Trials in which no nose-pose occurred within 3 min were excluded from analysis. Percentage correct response was counted as the number of correct responses over the number of total nose poke trials. Light stimulation was given only during the trials, but not during the inter-trial intervals.

Dissimilar odor discrimination continued for minimum 7 days until rats reached two consecutive days of 80% success rate, followed by similar odor discrimination for 10 days.

##### Odor Valence Tests

For conditioned odor preference test (COPT), rats underwent a single 30 min session of habituation in a t-maze (long arm 183 cm × 19 cm, neutral arm 19 cm × 19 cm; 20.5 cm high wall) on day 1. On day 2, O1 and O2 sponges were placed in two arms, with positions counter-balanced between morning and afternoon sessions. Time spent in the corresponding arms for O1 and O2 during 10 min session was recorded. On day 3, rats were confined to O1 arm for 10 min in the morning with light stimulation; and to O2 arm in the afternoon without light stimulation. The odors were switched in the arms on day 4 and the same conditioning as day 3 was repeated. Rats were trained for 1 trial or 3 trials (repeating procedures as in day 4-5 three times) and data were pooled. On the testing day, arm time was measured in morning and afternoon sessions with O1 and O2 sponges in exact same manner as in day 2.

For real-time odor preference test (ROPT), 122 cm long arm was used. Day 1 habituation and day 2 baseline light response were conducted in the same manner as in COPT. On day 3, rats explored the maze freely for two 10 min sessions in the morning and in the afternoon. Light stimulation started upon rat entering the O1 arm and stopped upon rat leaving the O1 arm. Odor positions were switched between morning and afternoon sessions.

##### CFos and Npas4 induction experiments

For cFos induction in Fig. 4&6, rats were habituated to experimental environment for two days. On third and fourth days, rats were optically stimulated with either phasic or tonic patterns in their home cages, for 10 min/day, while being exposed to an odorized sponge (benzaldehyde 0.005%). Ninety min following the stimulation on fourth day, rats were anesthetized, perfused and brains were collected.

For Npas4 induction to validate light activation in Fig. 1, rats were habituated to the experimental environment for one day. Next day they were perfused 60 min following light stimulation in their home cage.

### Immunohistochemistry and Histology

Rats underwent trans-cardiac perfusion with cold isotonic saline followed by 4% paraformaldehyde. The brains were extracted and kept in 4% paraformaldehyde. Brains were then sectioned in a vibratome (Leica VT 1000P; Leica Biosystems, Ontario, Canada) in 50 μm thick coronal slices and saved in polyvinylpyrrolidone solution. For immunohistochemistry with Npas4 and DBH, slices were washed in PBS, and then incubated with primary antibodies.

Npas4 (1:500, Thermo Fisher Scientific, Waltham, MA) primary antibody mixed in phosphate buffer saline (PBS) with 2% normal goat serum and 0.2% Triton-X was applied for three nights at 4°C. Then following 3 × 10 min wash in PBS, biotinylated anti-rabbit secondary antibody (1:1000; Vector laboratories, Burlingame, CA) was applied. After 2 hrs of secondary incubation and 3 × 10 min wash, slices were incubated in avidin-biotin complex for 1.5 hrs followed by PBS wash. Slices were then placed in a solution containing 15 μl SG Grey (Vector laboratories, Burlingame, CA) chromogen with 24 μl peroxide per ml of PBS. After optimum color development, slices were washed in PBS, dried overnight, dehydrated in graded ethanol and mounted with permount.

For DBH staining, after primary antibody (1:500, EMD Millipore, Burlington, MA) incubation, slices were incubated with fluorescence conjugated anti-mouse secondary antibody (1:1000; Invitrogen/Thermo Fisher Scientific, Waltham, MA) in room temperature for 2 hrs, followed by cover-slipping with Vectashield® antifade mounting medium (Vector laboratories, Burlingame, CA).

For cFos and TH co-labeling, 50 μm thick free floating sections from phasic and tonic stimulated rats, belonging to similar positions in antero-posterior axis of the brain as determined by unaided visual observation, were chosen in an unbiased manner. The sections were washed in Tris buffer (0.1M, pH 7.6) twice for 10 min each. Then 10 min in Tris A (0.1%TritonX in Tris buffer) and Tris B (0.1% TritonX and 0.005% BSA in Tris buffer) before applying blocking solution of 10% normal goat serum (Sigma-Aldrich, Oakville, Canada) for 1 hour. This was followed by 10 min wash each in Tris A and Tris B before incubating in 1:2000 primary antibody solution prepared in Tris B at 4°C (TH, EMD Millipore, Burlington, MA; cFos, Cell Signaling, Danvers, MA). After two nights, sections were washed for 10 min each in Tris A and Tris B and incubated in 1:1000 secondary antibody solution prepared in Tris B at 4°C (anti-rabbit Alexa 647, anti-mouse Alexa 488; Invitrogen/Thermo Fisher Scientific, Waltham, MA). This was followed by 10 min wash each in Tris A, Tris D (0.1%Triton X and 0.005% BSA in 0.5M Tris buffer) and Tris buffer respectively. Finally, sections were mounted with antifade mounting medium.

Nissl staining was done by rehydrating the slides in graded ethanol, incubating in 0.5% cresyl violet for 8 min, washing in distilled water for 1 min and then dehydrating in graded ethanol. After a 5-min xylene step, slides were coverslipped with permount.

### Image acquisition and analysis

Images were acquired by Aptome2 (Zeiss, Toronto, Canada), EVOS 5000 (Thermo Fisher Scientific, Waltham, MA) and BX-51 (Olympus, Richmond Hill, Canada) for fluorescent and bright-field images. Images were acquired similarly for phasic and tonic stimulated rats, keeping gain and exposure time the same throughout each experiment. Images were analyzed by ImageJ software. Images underwent background subtraction before manual cell counting. For LC activation success, the number of Npas4^+^ cells was counted. For BLA and VTA activation, cFos^+^ cells were counted. Three to six images per animal were analyzed and values from both hemispheres were averaged. A subset of the images was analyzed blindly.

### Statistical Analysis

One-way repeated ANOVAs were used in Fig. 1C to compare the frequency changes pre-, during, and post-light stimulation. A 3 × 5 mixed ANOVA was used in Fig. 1D to compare the effects of different light activation frequencies on LC firing. Two group comparison in Fig. 1F was subjected to Student’s t-tests (unpaired, two-tail). Two-way mixed ANOVAs were used to compare the effects of different patterns of light on general behavior in Fig. 2, followed by One-way repeated ANOVAs for each group, and *post-hoc* Bonferroni tests among light patterns or t-test between the control and ChR2 groups. Two-way mixed ANOVAs followed by linear trend analyses were used to determine statistical significance for the odor discrimination experiments, followed by t-tests between the control and ChR2 groups in Fig. 3 and Fig. 4. T-tests were used in Fig. 4 to compare cell numbers and percentage activation. A mixed ANOVA was used to compare the two odor valences in the control, ChR2 groups and NE blockade groups in ROPT and COPT in Fig. 5, followed by one-way ANOVAs on each group and *post-hoc* Bonferroni tests. T-test was used in two-group comparisons in Fig. 6. Data are presented as Mean ± S.E.M in the graphs.

## ACKNOWLEDGEMENT

This work was supported by Natural Sciences and Engineering Research Council of Canada discovery grants to Xihua Chen (#261384-2008) and Qi Yuan (RGPIN-2018-04401). The authors wish to thank Dr Karl Deisseroth’ laboratory at Stanford University for support and providing AAVs; Special thanks to Ms. Charu Ramakrishnan.

