## Supplementary figures and images for "Locus coeruleus patterns differentially modulate learning and valence in rat *via* the ventral tegmental area and basolateral amygdala respectively"

### Supplementary Figures

**SUPPLEMENATRY FIGURES**
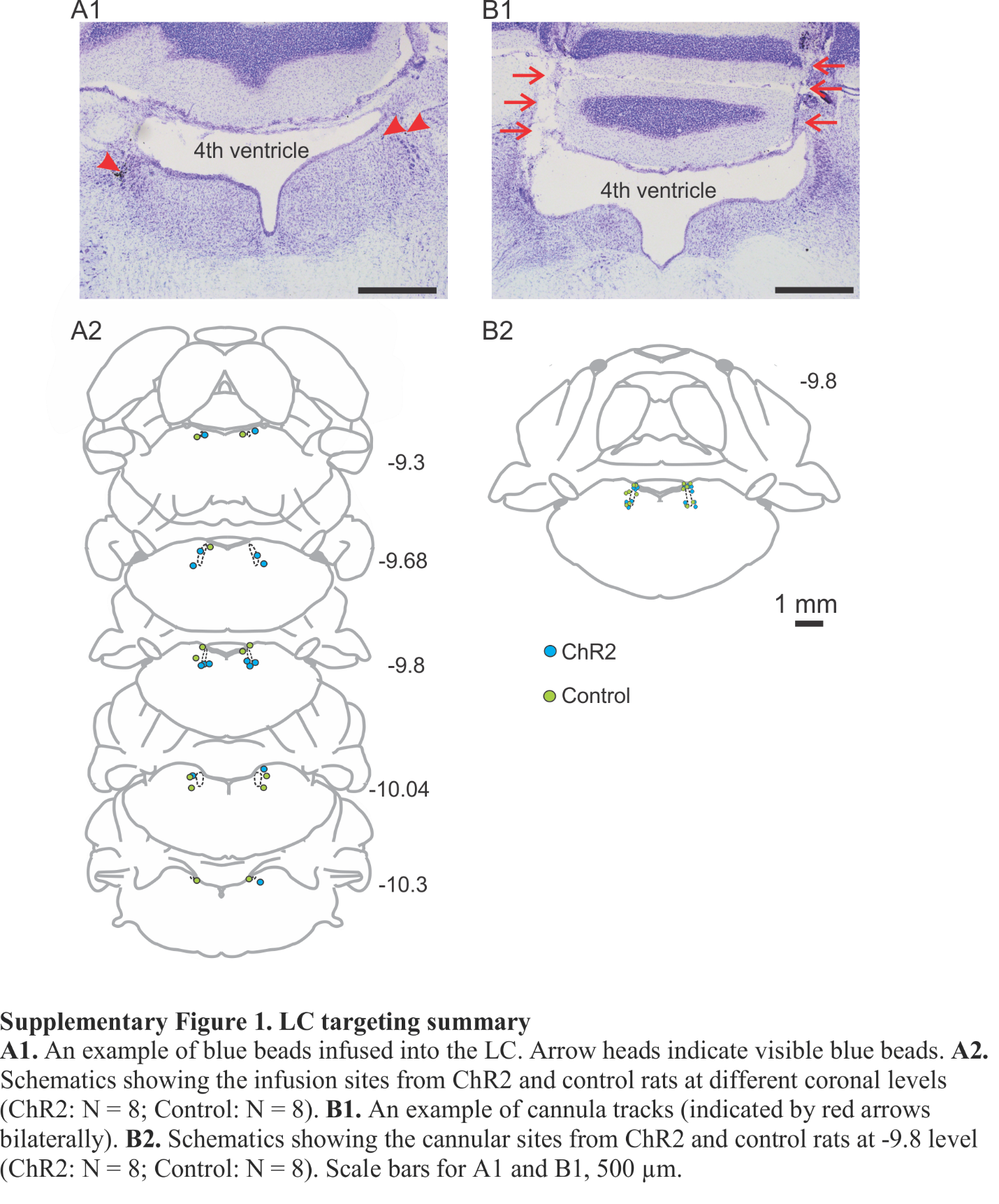

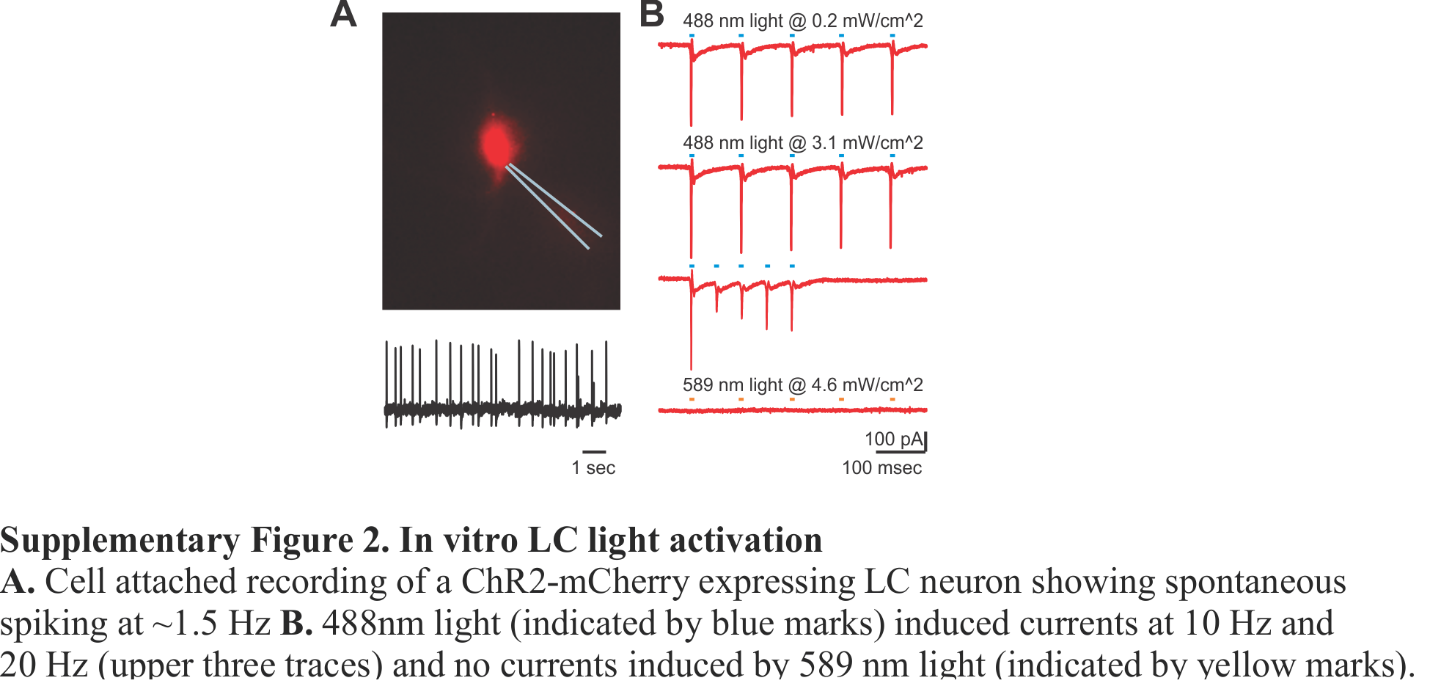

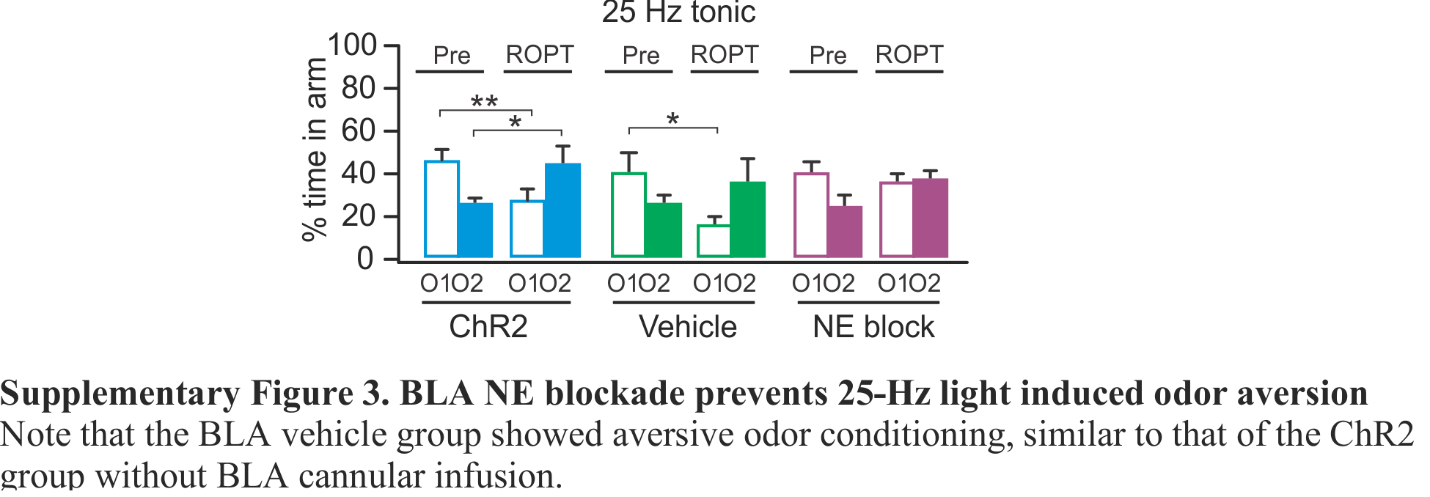
